# Indirect effects on fitness between individuals that have never met via an extended phenotype

**DOI:** 10.1101/442335

**Authors:** David N. Fisher, Jessica A. Haines, Stan Boutin, Ben Dantzer, Jeffrey E. Lane, David W. Coltman, Andrew G. McAdam

## Abstract

Interactions between organisms are ubiquitous and have important consequences for phenotypes and fitness. Individuals can even influence those they never meet, if they have extended phenotypes which mean the environments others experience are altered. North American red squirrels (*Tamiasciurus hudsonicus*) guard food hoards, an extended phenotype that typically outlives the individual and is almost always inherited by non relatives. Hoarding by previous owners can therefore influence subsequent owners. We found that red squirrels bred earlier and had higher lifetime fitness if the previous owner was a male. This was driven by hoarding behaviour, as males and mid-aged squirrels had the largest hoards, and these effects persisted across owners, such that if the previous owner was male or died in mid-age subsequent occupants had larger hoards. Individuals can, therefore, influence each other’s resource dependent traits and fitness without meeting via extended phenotypes, and so the past can influence contemporary population dynamics.

## Introduction

Organisms socially interact when they mate, fight and compete for resources. Social interactions are thus ubiquitous throughout the natural world (Frank 2007). Social interactions mean individuals will influence the traits and fitness of other individuals (Scott 1977; Moore *et al.* 1997). For instance, individuals can behave cooperatively through behaviours such as pack-hunting or cooperatively raising young, which can have positive effects on fitness components of others. Alternatively, individuals can compete for limited resources, which negatively influences the fitness of others. Therefore, an organisms’ fitness can be determined not just by its own traits (and genes), but also by the traits of others with which it interacts (Okasha 2004a, b). This potentially complicates the relationship between an organism’s traits and its fitness, altering the evolutionary response to selection. Additionally, if these social effects have a heritable component, then phenotypes within a population are also influenced by genes of other individuals. These are referred to as indirect genetic effects (IGEs; Griffing 1967; Moore *et al.* 1997; Wolf *et al.* 1998) in contrast to more traditional direct genetic effects (DGEs, an individual’s own genes influencing its phenotype and so fitness). IGEs can fundamentally alter evolutionary trajectories (Bijma & Wade 2008), so social interactions are predicted to play an important role in evolution (Bailey *et al.* 2018).

It is often implicitly assumed that organisms need to meet to influence each other, but this need not always be the case. If organisms possess extended phenotypes (traits of the individual that exist outside their physical body; Dawkins 1978, 1982) then they can alter the environment that others experience. “Ecosystem engineers” (Jones *et al.* 1994) such as earthworms (Thompson *et al.* 1993) and beavers (*Castor spp.*) (Naiman *et al.* 1986; Rosell *et al.* 2005) are well known to alter the environment for their own benefit (see also: “niche construction”, Odling-Smee *et al.* 2003; Scott-Phillips *et al.* 2014), and so will influence any organisms that use the modified environment. By altering the environment that another individual might experience, organisms can influence the plastic traits of others, or perhaps even their fitness, despite never actually meeting (Laidre 2012). Yet how important such indirect effects are outside of these classic “ecosystem engineers” is not clear.

Like any other trait, an extended phenotype could be under selection, and so might be expected to evolve (Dawkins 1982). However, predicting the microevolution of extended phenotypes requires estimates of the genetic variances underpinning them, yet such estimates are rare (Saltz & Nuzhdin 2014). An exception is the case of oldfield mice (*Peromyscus polionotus*), which have extended phenotypes in the form of long entrance tunnels to their burrows, from which there may be a second “escape tunnel” (Weber *et al.* 2013). A low number (possibly as few as three) loci are thought to control burrowing behaviour, specifically the differences in tunnel length and the tendency to build escape tunnels compared to a sister species (*P. maniculatus*) (Weber *et al.* 2013), with 8.2% of phenotypic variance in adult tunnel length attributed to a single nucleotide polymorphism (Metz *et al.* 2017). Yet the generality of such findings for behaviours expressed in the wild, and so whether extended phenotypes are typically heritable, is not known. If individuals affect each other through extended phenotypes, and the extended phenotype possesses genetic variance, part of this indirect effect could be genetic, and so be an IGE. However, whether genetic variance in extended phenotypes means IGEs for other individuals experiencing the modified environment has not been considered (in general, estimates of IGEs in natural populations are rare, but see: McAdam & Boutin 2004; Brommer & Rattiste 2008; Wilson *et al.* 2011; McFarlane *et al.* 2015).

It is, therefore, necessary to test how extended phenotypes can cause individuals to influence the phenotypes and fitness of those they never meet, and to determine the quantitative genetic underpinnings of extended phenotypes. To do this, we studied a population of North American red squirrels (*Tamiasciurus hudsonicus*; hereafter “red squirrels”) at a study site in the Yukon Territory of Canada for 30 years. Red squirrels defend territories of around 0.34 ha (LaMontagne *et al.* 2013), centred on a “midden”, a pile of white spruce (*Picea glauca*) cone scales, within which red squirrels cache food they require to survive over winter (Smith 1968b). Stored food typically consists of white spruce cones (the red squirrels’ main food source in this population) harvested that autumn (“new cones”) as well as cones stored from previous years’ crops (“old cones”). Middens vary in the number of cones of each type they contain, with larger hoards leading to improved overwinter survival of the occupant (Larivée *et al.* 2010; LaMontagne *et al.* 2013), earlier breeding in the spring for females and increased reproductive success for males (Haines 2017; Haines *et al., in sub.*).

Along with being an important contributor to the current occupant’s survival and fitness, middens last well beyond the lifespan of a single individual. Any given midden may be defended over time by a sequence of many individuals that are typically not related and do not overlap in tenure (Hatt 1929; Smith 1968b, 1981), although in some cases a female squirrel will “bequeath” her territory to one of her offspring, and leave to find another territory (Price & Boutin 1993; Berteaux & Boutin 2000; Boutin *et al.* 2000; Lane *et al.* 2015). Spruce cones cached in a midden can be consumed at least four years post-caching (Donald & Boutin 2011; S. Boutin, unpublished data), so cones cached by previous owners can be used by the inheritor of a territory. Therefore, the number of cones cached in a squirrel’s midden is an extended phenotype, and furthermore may well be influenced by both the current and the previous occupant of the territory (Fig. 1). This then creates a mechanism through which the previous owner can influence the resource-dependent traits, and so possibly fitness, of the current occupant. In addition, if there is genetic variance in current hoard size, there could be IGEs influencing the phenotype of the current owner through the extended phenotype of the previous owner.

**Figure 1.**
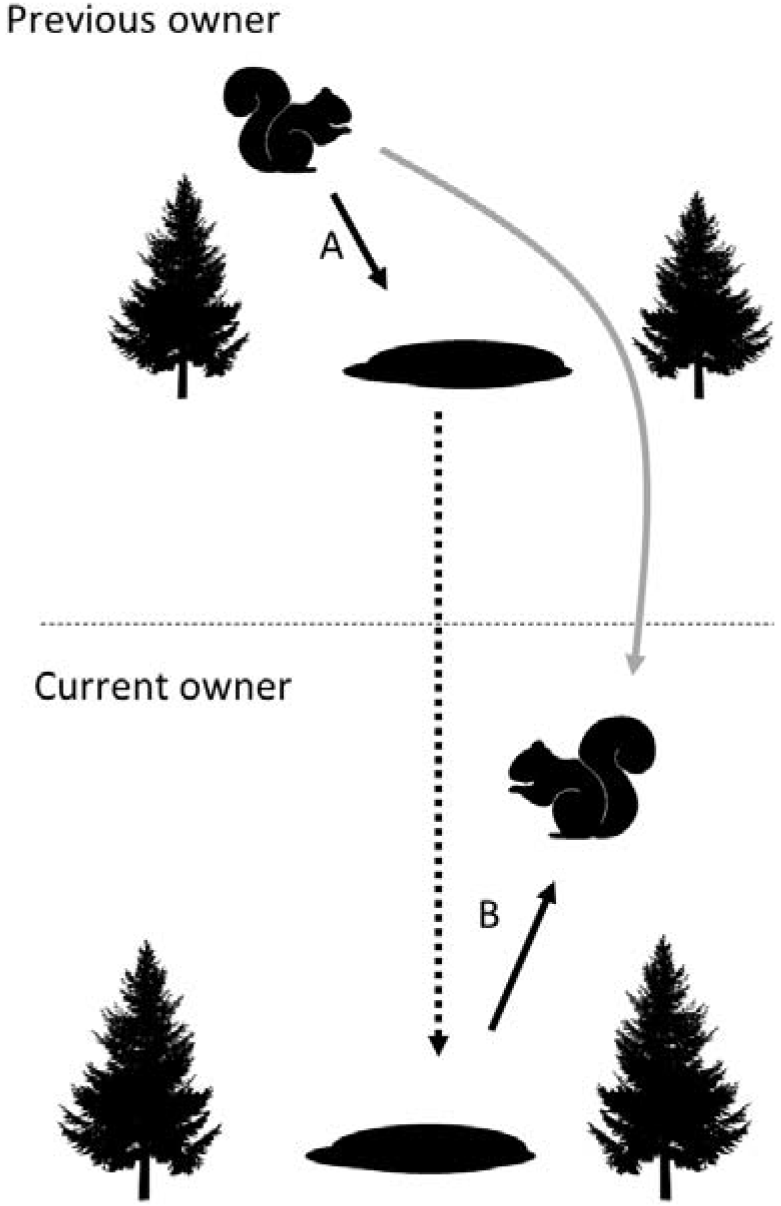
Extended phenotypes, in the form of a resource cache, can cause previous territory occupants to influence the current occupant. A squirrel can influence the size of its resource cache (line labelled A). Some of this alteration may then retained beyond the life of the current occupant, through to the next occupant (dashed line) and this variation then may influence a resource dependent trait of the new occupant (line labelled B). This implies that an individual influences the traits of another individual (grey line), which is typically only considered when organisms directly interact. Here however it may occur despite the individuals never being alive at the same time, through an extended phenotype.

We identified how the previous occupant of the territory influences the traits (cone hoard sizes, as well as the date a female gives birth in the spring; “parturition date”), and the fitness (estimated as lifetime reproductive success; LRS) of the current owner. Additionally, to assess whether cone hoards are heritable, we used an “animal model” (Kruuk 2004; Wilson *et al.* 2010) based on a multigenerational pedigree for this population to estimate DGEs of the current owner on the hoard sizes of old and new cones and IGEs from the previous territory occupant on cone hoards, as well as for parturition date and LRS. Since they are placed in the ground in the autumn when hoard size is measured, we expected no influence of the previous owner on the size of new cone hoards, but we did expect the previous owner to influence the size of old cone hoards. Females give birth earlier when they have access to increased food resources (Réale *et al.* 2003b; Kerr *et al.* 2007), so we expected that traits of the previous occupant that were associated with larger cone hoards would be the traits that cause the subsequent hoard owner to breed earlier. We made the same prediction for LRS; traits of the previous owner that are associated with larger hoard sizes should confer higher fitness on the current occupant of the territory.

## Methods

### Data collection

We followed individually marked squirrels as part of the Kluane Red Squirrel Project in the Yukon Territory of Canada between 1987 and 2017. We used different subsets of this entire dataset for our analyses here. In each year, we monitored marked females in two unmanipulated 40-ha. study areas (“Kloo” and “Sulphur”) for signs of pregnancy and to tag their pups. We also tagged any immigrants and, therefore, tracked all squirrels resident in the population for their entire lifetime (see: McAdam et al. 2007 for further details on the study system). We enumerated the entire population in spring and autumn censuses to determine ownership of territories. These territories are exclusive and are based around large piles of discarded cone scales (the midden). Middens are semi-permanent, so the same midden can be owned sequentially by multiple different squirrels across an extended period of time. For example, one midden remained active throughout the entire study period (between 1987 and 2017) and was owned by 13 different squirrels across this 31-year period. We assigned ownership of a midden based on territorial vocalizations called ‘rattles’ (Smith 1968a, 1978), and only included squirrels owning a midden in our analysis, i.e. those who had recruited into the population. We focus here on territory ownership in the autumn, as this is when spruce cones are available to be stored, and so when we measured the size of cone hoards in middens. We recognised previous territory owners when the identity of the territory owner in one autumn was different to that in the previous autumn. We only included the first instance of a squirrel holding a given territory in the analysis, so there were not multiple records if a squirrel held the same territory for several years, but we did include a squirrel multiple times if it was observed holding different territories in different years. Each squirrel-midden combination, therefore, only had a single previous owner, and since middens left undefended through the death of the owner are typically taken over by new individuals quickly (Price *et al.* 1986; Siracusa *et al.* 2017a, b), the previous owner typically held the territory a single year ago. If the midden was undefended for >1 year, the previous owner would be more than 1 year in the past, which we accounted for by fitting an interaction between the traits of the previous owner and the number of years between them and the current owner (see Data analysis).

Between 2012 and 2017 we estimated the number of cones stored in the primary middens of all squirrels defending territories. We did this in late September after most caching was completed. Some squirrels owned more than one midden in the autumn, but we did not consider secondary middens as they are not held by the majority of squirrels and are not used extensively for resource storage when they are held. For each midden, we identified its perimeter as the location where the cone bracts gave way to typical forest floor, measured the longest axis and the perpendicular width at the midpoint of the long axis. Assuming an elliptical shape (following: Larivée *et al.* 2010), we estimated the area as :

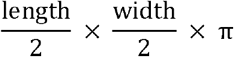

Within each quarter of the midden, we placed two 30cm X 30cm square quadrats at approximately ¼ and ¾ of the distance from the centre of the midden towards the perimeter, such that each quarter of the midden had two quadrats placed wholly within it, giving eight in total (method adapted from Larivée et al. 2010). If the midden was very small, such that a quarter could not fit two quadrats, then only a single quadrat was placed. We then excavated all quadrats to a depth of 10cm from the surface of the bract pile and counted the number of old and new cones found within. If an excavated tunnel was found, new and old cones inside were counted to a depth of 30cm, but not out of the area of the quadrat. If cones are not stored in middens they will open, releasing their seeds. This makes them useless as food to red squirrels, so we did not count opened cones. We calculated the number of old and new cones per cm^2^ of quadrat (accounting for the number of quadrats), multiplied by the area of the midden, and then rounded to the nearest whole number, to give an estimate of the size of the old and new cone hoard in the midden. We were able to separately count old and new cones, as new cones possess hues of purple and green, and were often sticky with sap, while old cones were dark brown and never sticky.

For the years outlined above, we were able to assess the effect of previous owners on old and new cone hoard sizes. Across a much-extended period of data collection, we were also able to estimate effects of previous owners on female parturition dates (springs of 1992-2017) and the LRS of males and females (squirrels born from 1991-2009; although LRS of males was not available until 2003, see below). Due to maternities identified at birth since 1987, and paternities identified based on genetic analysis of ear biopsies since 2003, we have very good information on the relatedness among individuals within our study areas, giving a well-resolved pedigree (Gunn *et al.* 2005; Lane *et al.* 2007; McFarlane *et al.* 2011). We estimated LRS as the total number of pups an individual dammed or sired that survived to 200 days, i.e. had survived their first winter and recruited into the population as adults. We excluded individuals for which we had missed one or more breeding events, individuals who died of unnatural causes such as dying in a trap, individuals who were born after 2009 (as their LRS would be underestimated), as well as males pre-2003 for which we were not able to asses siring success. As mentioned above, female red squirrels may bequeath their territory to an offspring, which tends to be a daughter and the fastest growing pup in the litter (Berteaux and Boutin 2000; Robertson *et al., in prep*). We repeated all analyses with all instances of the previous occupant being the mother excluded. Those results are qualitatively similar to the ones we present here, and so we present them in the supplementary materials.

### Data analysis

We used variations of the animal model (Kruuk 2004; Wilson *et al.* 2010) to determine (1) effects of the previous occupant on current cone hoard estimates, parturition dates and LRS and (2) direct and indirect genetic variance for cone hoard sizes, parturition dates, and LRS. This involved four linear models, either with the old cone hoard size, new cone hoard size, parturition date or LRS as the response variable, all fitted in R using the package “MCMCglmm” (Hadfield 2010). We also used the pedigree to calculate the genetic relatedness between the territory owner and the previous occupant (*r*).

For all models, we included an “animal” term, linked to the pedigree, to estimate the direct genetic variance for each trait. Heritabilities were then calculated using the package “QGglmm” (de Villemereuil *et al.* 2016; as estimates are initially on the latent scale, not the response scale), with the among-year variance excluded, as selection and therefore the relevant heritability measure is primarily within-years (Réale *et al.* 2003a; Lane *et al.* 2018). For cone hoards and parturition dates we also included the random effect of squirrel identity, since some squirrels had multiple measurements of hoard sizes and parturition dates. This allowed us to estimate permanent environment effects (where individuals are consistently different from one-another, but for reasons not due to additive genetic differences or other variables included in the model). In every model we also fitted the identity of the previous owner as an additive genetic effect, which allowed us to estimate variance in IGEs. We also estimated the direct-indirect genetic covariance, to determine whether genes for large cone hoards were associated with genes for leaving the next owner more or less cones (and so on for each trait analysed). We included in all models the random effect of year, to account for year-to year variations in the traits at the population level.

We added a different set of fixed effects to each model to control for contemporary factors (traits of the current occupant and environmental influences) that might influence the traits. To the models of old and new cone hoard sizes we included the current occupant’s sex, age, study area, and the linear effect of year. Within an individual’s lifetime cone hoards have been shown to increase and then decrease (Haines 2017). We therefore estimated a different relationship between hoard sizes and age before and after the approximate age of peak cone hoard sizes by including a term for whether the squirrel was older than three years or not (approximately the age of peak cone hoards), and the interaction between this term and age. Fitting two relationships like this is preferable to using age^2^ to detect initial increases followed by decreases (Simonsohn 2018; e.g. due to senescence) although the results for models in which age was fitted as a quadratic term instead led to similar conclusions (not shown). For parturition date, we included the occupant’s age in years as a categorical variable (with 7- and 8-year olds, the oldest, grouped together), to account for non-linear trends (typically late parturition dates as a yearling breeder and potential senescent declines in old age), the individual’s study area, and a linear effect of year. For LRS we included the individual’s study grid, a linear effect of year, and whether the individual experienced a “mast year” (when spruce trees produce a super abundance of cones; Kelly 1994; Lamontagne & Boutin 2007) or not in their lifetime, as this has been shown to greatly increase LRS (Descamps *et al.* 2008; Hämäläinen *et al.* 2017).

To investigate whether the previous occupant influenced the traits of the current owner, in all models we included the sex and lifespan of the previous owner as fixed effects, with separate relationships for lifespan of the previous owner below and above three years of age, in the same manner and for the same reasons as above. We included sex of the previous owner because males have been found to cache more and have larger hoards (Donald & Boutin 2011; Archibald *et al.* 2013), which could influence the next owner of the territory. We also included the interaction between these traits of the previous owner and the time in years between the current and previous owners, because we predicted that previous occupants farther in the past should affect the current occupant less. We also included the main effect of this time span, which represented the length of time the midden was unoccupied before the current occupant. As unoccupied middens will have their stored cones gradually removed by other red squirrels, we expected larger values of this time span to be associated with smaller old cone hoards, later parturition dates, and lower LRS estimates.

For cone hoards and LRS we used a Poisson error-structure with a log-link function, an inverse Gamma prior for each random effect (V = 1, nu = 0.002, chosen to be non-informative), and 500,000 iterations, with the first 10,000 discarded, and 1/40 of the remaining iterations kept, to form the posterior distributions. For parturition date we used the same specifications, except a normal error structure and link function. For all models, we standardised each continuous predictor variable by subtracting the mean from each value and then dividing by the standard deviation, which improves model convergence and interpretability of regression coefficients (Schielzeth 2010). We also scaled parturition dates in this way, but not cone hoard sizes or LRS, as this transformation gives negatives and non-integers, unsuitable for Poisson models. Importance of terms was judged by the distance of the mode of the posterior distribution from zero, and the spread of the 95% credible intervals. Successful convergence was assessed with Heidelberger and Welch’s convergence diagnostic (Heidelberger & Welch 1983), while three runs of each model performed to ensure different chains reached the same qualitative result.

## Results

### Phenotypic effects

The sex of the previous midden owner had important consequences for the current midden owner, which acted through the extended phenotype of the size of the hoard. Males hoarded more new cones, (coefficient plots for all models are given in Fig. 2, while sample sizes and other summary information are given in Table 1). This effect on hoard size carried over to the next owner of the midden. If the previous midden owner was a male, the current occupant had a larger old cone hoard (Figs. 2a & 3a), an earlier parturition date (Figs. 2c & 3b, although this effect marginally overlapped zero), and a higher LRS (Figs. 2d, 3c). Given a female squirrel of average age (and so not post-peak) on the Kloo study area, with a previous owner also of average age and not post-peak, the previous owner being male rather than female would result in an extra 1318 old cones in the hoard of the current occupant, would mean the female gave birth 2.26 days earlier, and, assuming the female did not experience a mast, means she would have 1.79 instead of 1.2 offspring survive to 200 days over her lifetime. As expected, this effect of the sex of the previous occupant on the size of the old cone hoard weakened with increasing number of years between the current and previous owner (Fig. 2a). Surprisingly, the effect of whether the previous owner was a male on LRS was enhanced with increasing time between the previous and current owner (although this effect marginally overlapped zero; Fig 2d).

We also found that the lifespan of the previous midden owner influences the current midden owner through hoard size. Within the lifetime of current owners, old and new cone hoard sizes both showed no initial increase before three years of age, and then declined with age for squirrels that were older than three years (Fig. 4a & b). Concurrent with this late-life decline, red squirrels who lived longer than three years of age left fewer old cones to the next occupant of the territory the longer they lived (Fig. 4c). We also found that parturition dates were later if the previous occupant lived longer, with no change in this relationship pre- and post-three years of age (Fig. 4d). Note however that each of these effects slightly overlapped zero, suggesting they should be interpreted with caution. Parturition dates were also latest in yearlings, earliest at ages 5 and 6, and tended to be later at ages 7 and 8. LRS was unaffected by the age of the previous owner.

**Figure 2.**
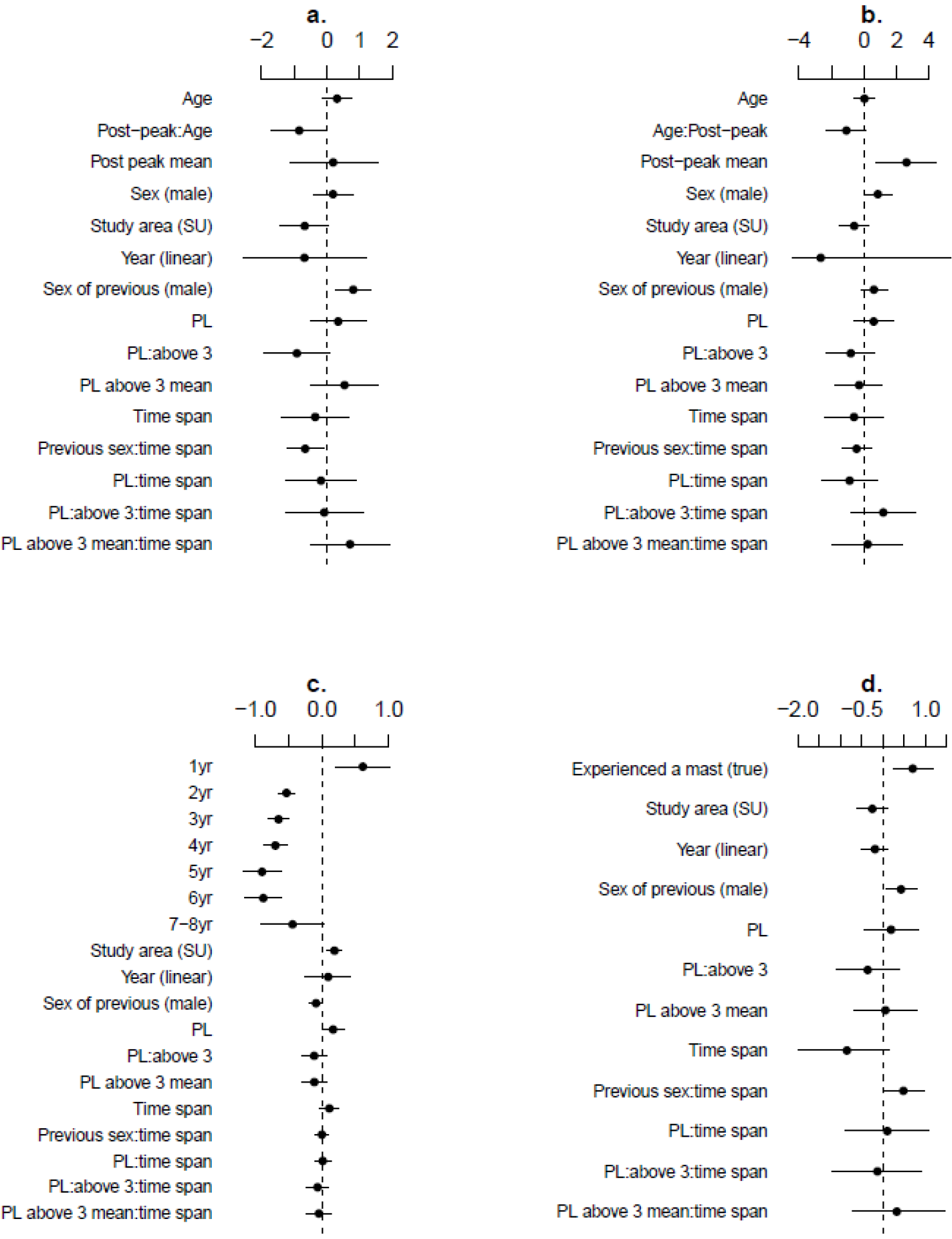
Coefficient plots displaying estimated means and 95% credible intervals (CRIs) of the models for a) old cone hoards, b) new cone hoards, c) parturition dates (note negative values indicate earlier dates) & d) lifetime reproductive success. “Age” refers to the effect of age of the current occupant on the trait, while “Post-peak:Age” refers to the additional effect of age when the individual was older than 3 years. “Post-peak mean” indicates whether traits in individuals older than 3 had different means. “Previous sex” indicates the additional effect of the previous owner being a male, with it being a female the default. “PL” refers to the previous occupant’s lifespan, while “PL:above 3” refers to the additional effect of this lifespan if it was above 3 years. “PL above 3 mean” indicates whether traits in individuals where the previous territory occupant lived longer than 3 years had different means. “Time span” is the time period between the previous and current occupant; it is interacted with each trait of the previous occupant (e.g. “Previous sex:time span”). For new cone hoards the x-axis has been truncated to display the effects nearer zero; the CRIs for the effect of year were −13.8 to 9.3.

**Figure 3.**
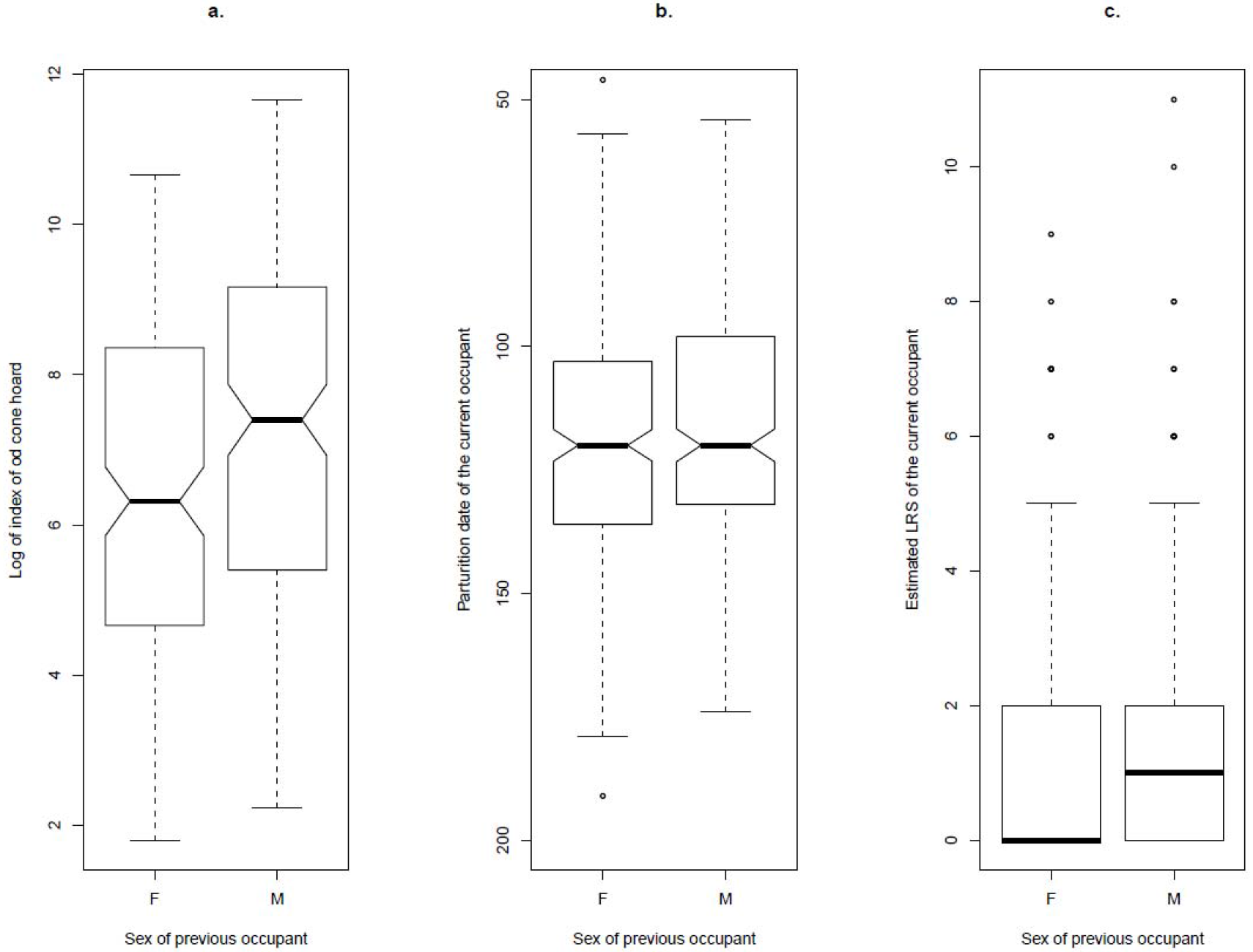
a) Old cone hoards were larger when the previous occupant was a male. Old cone hoards have been ln-transformed to improve viewability. b) Parturition dates (days since 1^st^ January) tended to be earlier when the previous owner was a male. c) Estimated LRS was higher when the previous owner was a male. Notches in a & b indicate 95% confidence intervals of the median (the thick black bar in all plots).

**Table 1.**
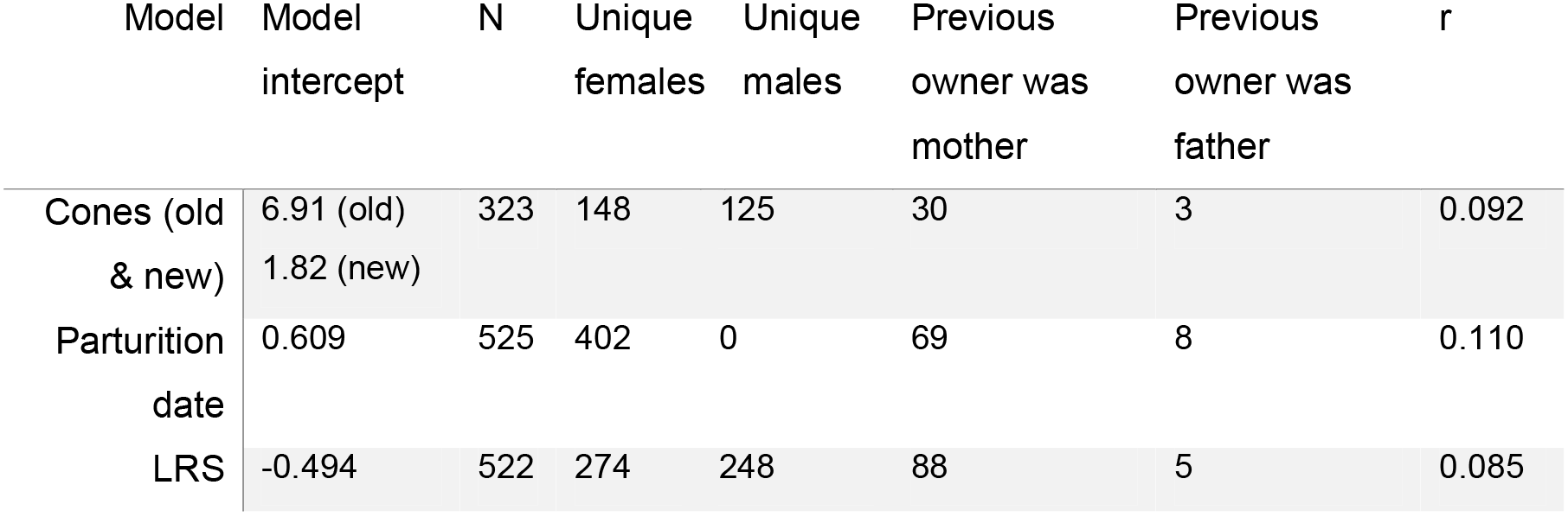
Model intercepts, number of measures (N), number of unique females and males, number of instances the previous owner was the mother or the father, and the mean coefficient of relatedness (*r*) between current and previous owners, for each of the four analyses (note the dataset for old and new cone hoards is the same).

| | Model intercept | N | Unique females | Unique males | Previous owner was mother | Previous owner was father | $r$ |
| --- | --- | --- | --- | --- | --- | --- | --- |
| Cones (old & new) | 6.91 (old)<br>1.82 (new) | 323 | 148 | 125 | 30 | 3 | 0.092 |
| Parturition date | 0.609 | 525 | 402 | 0 | 69 | 8 | 0.110 |
| LRS | -0.494 | 522 | 274 | 248 | 88 | 5 | 0.085 |

**Figure 4.**
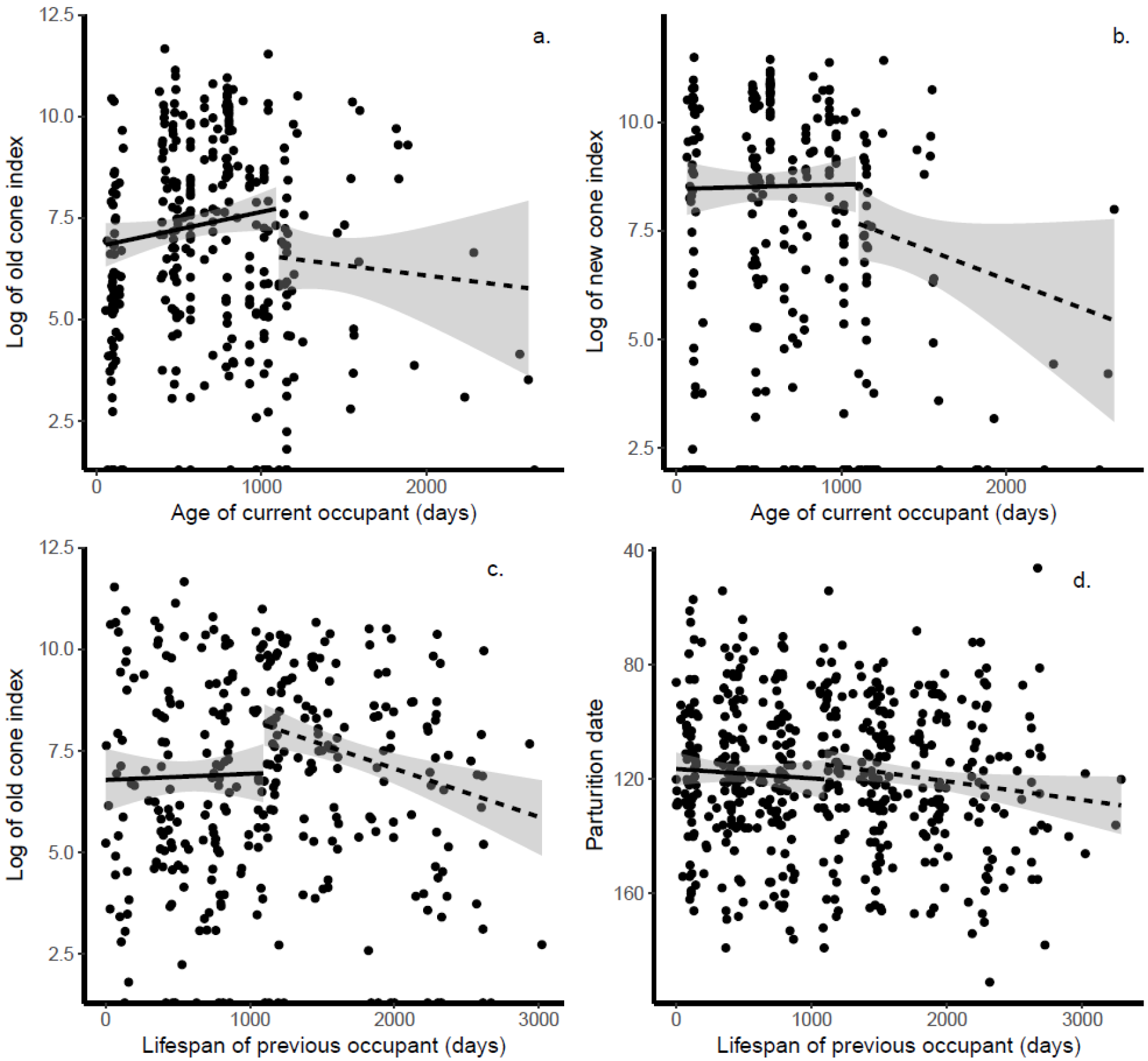
a) Old cone hoards declined with age in squirrels older than three years. b) new cone hoards declined with age in squirrels older than three years. c) Cone hoards were largest if inherited from an individual of intermediate lifespan and declined if the previous owner lived beyond three years. d) parturition dates were later the long the previous individual lived. For all plots cone hoards have been logged to improve viewability.

While not of direct interest to this study, contemporary effects on traits remained. Individuals had a higher LRS if they experienced a mast year in their lifetime. Old but not new cone hoards were smaller and parturition dates later on one study area (Sulphur) compared to the other (Kloo), but LRS was not different. No trait showed a consistent change across years, while LRS was lower the longer the midden had been unoccupied (although this effect marginally overlapped zero), but this time period did not affect the other traits.

### Genetic effects

We found very low levels of genetic variance in old cone hoards and new cone hoards (estimates given in Table 2). These gave *h*^*2*^ estimates of < 0.001 for both types of cone. It is therefore unsurprising that there were no IGEs from the previous owner for any trait, although for old cone hoard the estimate mode was quite far from zero. The direct and indirect genetic effects did not significantly covary in any case, although for old cone hoards red squirrels with genes for large hoards tended to leave more cones to the next owner. Otherwise, parturition dates were heritable (h^2^ = 0.147), and LRS was less so (h^2^ = 0.099).

**Table 2.**
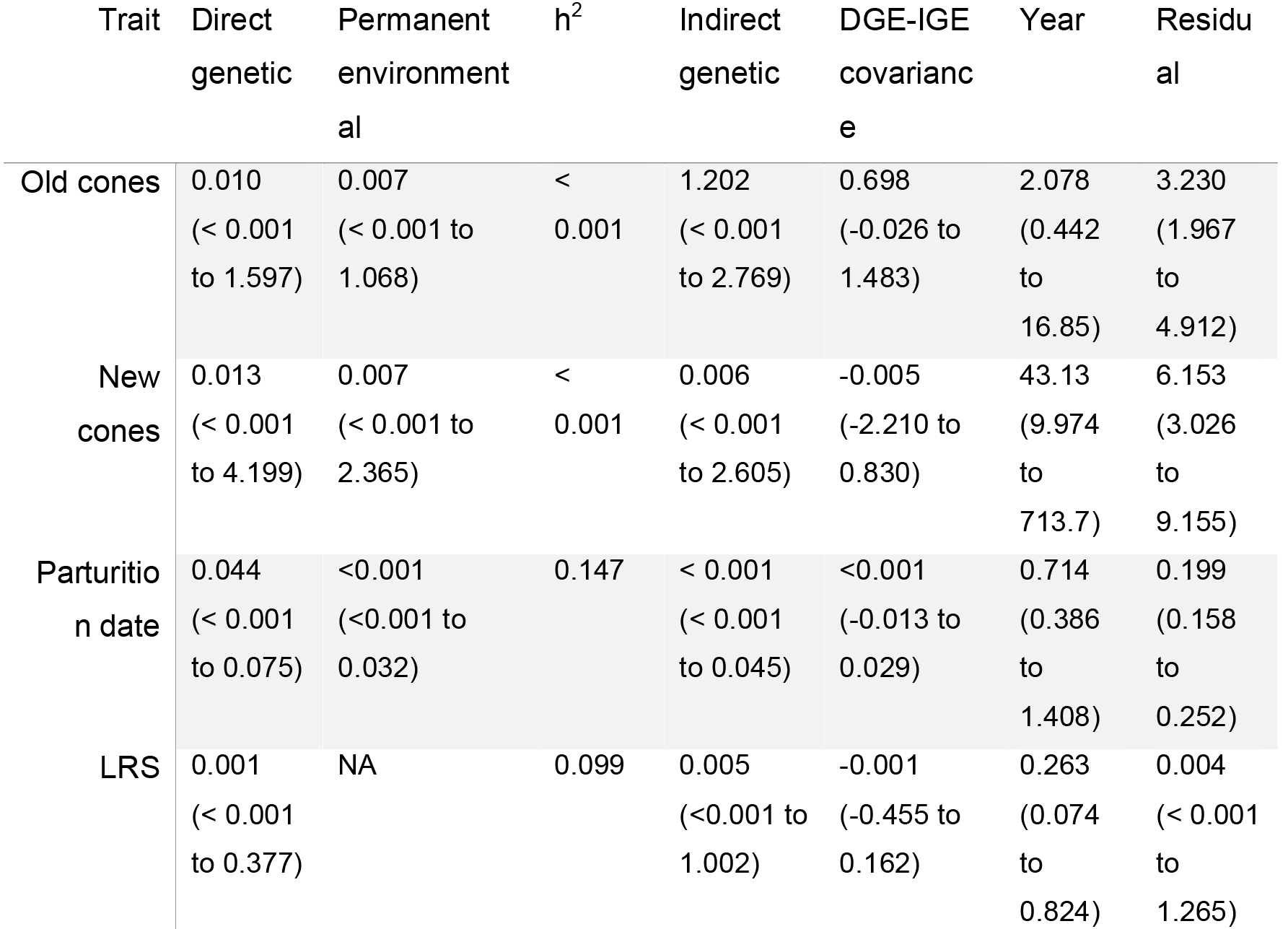
Variance component estimates for each model, as well as the point estimate for narrow-sense heritability (*h*^*2*^, calculated with the among-year variance excluded, see Methods). For variance components we give the posterior distribution mode, with the 95% credible intervals in parentheses. “NA” indicates the term was not estimated in the model. Note the “Year” term in the LRS model is birth year, whilst in the other models it is the year the trait was expressed.

| Trait | Direct genetic | Permanent environmental | $h^2$ | Indirect genetic | DGE-IGE covariance | Year | Residual |
| --- | --- | --- | --- | --- | --- | --- | --- |
| Old cones | 0.010<br>($< 0.001$ to 1.597) | 0.007<br>($< 0.001$ to 1.068) | $< 0.001$ | 1.202<br>($< 0.001$ to 2.769) | 0.698<br>(-0.026 to 1.483) | 2.078<br>(0.442 to 16.85) | 3.230<br>(1.967 to 4.912) |
| New cones | 0.013<br>($< 0.001$ to 4.199) | 0.007<br>($< 0.001$ to 2.365) | $< 0.001$ | 0.006<br>($< 0.001$ to 2.605) | -0.005<br>(-2.210 to 0.830) | 43.13<br>(9.974 to 713.7) | 6.153<br>(3.026 to 9.155) |
| Parturition date | 0.044<br>($< 0.001$ to 0.075) | $< 0.001$<br>($< 0.001$ to 0.032) | 0.147 | $< 0.001$<br>($< 0.001$ to 0.045) | $< 0.001$<br>(-0.013 to 0.029) | 0.714<br>(0.386 to 1.408) | 0.199<br>(0.158 to 0.252) |
| LRS | 0.001<br>($< 0.001$ to 0.377) | NA | 0.099 | 0.005<br>($< 0.001$ to 1.002) | -0.001<br>(-0.455 to 0.162) | 0.263<br>(0.074 to 0.824) | 0.004<br>($< 0.001$ to 1.265) |

## Discussion

Indirect effect have been suggested to play a “special role” in evolution (Bailey *et al.* 2018), as they cause individuals to influence each other’s phenotypes and fitness (Griffing 1967; Scott 1977; Moore *et al.* 1997). Here we have documented that indirect effects can occur between organisms that never meet and may not even have been alive at the same time, facilitated through an extended phenotype. Extended phenotypes have received much theoretical interest since Dawkins (1982) (Jones *et al.* 1994, 1996; Sterelny *et al.* 1996; Dawkins 2004; Jablonka 2004; Turner 2004; Bailey 2012), but extending empirical work to a range of systems has lagged behind. Many species hold territories that they may modify, for instance by digging out burrows, building nests or caching resources within them. Additionally, organisms may alter the environment by constructing some structure in it, such as a spider’s web (Blamires *et al.* 2017a, b). Therefore, there is the potential for effects mediated by these extended phenotypes to be widespread throughout the natural world, yet our study is the first to quantify influences on life-history traits and fitness of other individuals mediated through an extended phenotype.

### Sex of previous owner influences cone hoards, parturition dates and fitness

If the previous owner of the territory was a male, current occupants had earlier parturition dates and higher LRSs, due to inheriting larger hoards of old cone. Therefore, any trait which is related to how much an organism alters its environment might be expected to predict how much it influences other organisms indirectly. Often, only one sex of a species controls a territory, builds a nest, or engages in some form of environmental manipulation. It is therefore only this sex which is expected to exert indirect effects through these environmental changes, and anything which alters the survival and so prevalence of this sex, or its distribution in the environment, will alter the influence of these indirect effects.

Finding more cones in middens previously owned by males appears to suggest that males hoard more cones than females, then fail to use them, as they are still present for the next occupant. Male energy expenditure during the mating season is approximately as high as female energy expenditure during lactation (Lane *et al.* 2010), so it unlikely that males simply do not need to use these resources to fuel reproduction. Instead, males may hoard more as hoard size is related to the number of offspring they sire in the spring, which is not true for females (Haines 2017). Over-winter mortality would prevent males from consuming these additional cones during the mating season, leaving the larger hoard for the next owner. We also found that males did not have larger old cone stores, which might seem to contradict the general findings that they have larger cone hoards. However, this can be explained by the fact that that we used only the first instance of holding a territory for each squirrel, so males likely have not yet had time to enlarge their hoard of cold cones.

If male red squirrels leave behind more resources, it makes it beneficial for a juvenile squirrel to settle at a territory that had previously been held by a male, to take advantage of the extra resources. This is generally true for any case of environment manipulation: it would be advantageous for other organisms to detect and exploit any changes made by others. In this case, we do not believe there to be any choice in where a juvenile squirrel settles, as it tends to be close to the natal midden, and availability of territories is very limited, with competition for them intense (Price & Boutin 1993; Larsen & Boutin 1994; J. Robertson, unpublished data), especially in years of high density (Williams *et al.* 2014; Fisher *et al.* 2017). It may well be completely stochastic what sex the previous occupant of a territory was, although this may not be true in other systems. Ultimately, this effect of the previous owner will still have consequences for the fitness of the current occupant, and so a portion of the variance in reproductive success of a population can be attributed to the caching behaviour of now-dead individuals. The effects of the previous owner on fitness, therefore, have the potential to have interesting effects on the distribution of fitness in a population (discussed below).

### Largest hoards passed on by previous owners of intermediate age

We found that cone hoard sizes decreased beyond three years of age. This suggests that the ability to find and/or cache cones declines in old age and is perhaps an example of a senescent decline. Phenotypic senescence is increasingly commonly found in natural populations (Nussey *et al.* 2013), and has been detected in this study system before (McAdam *et al.* 2007; Descamps *et al.* 2008), including for cone hoard size (Haines 2017). This senescence in the stored resources an individual has access to is analogous to senescence in fat reserves in organisms that store energy on their bodies (“capital breeders”; Jönsson 1997; e.g. in sockeye salmon, *Oncorhynchus nerka;* Hendry & Berg 1999). This demonstrates senescence is a general phenomenon that even extends beyond the commonly considered case of “performance” traits such as body mass or clutch size, to include an extended phenotype, and so should be considered in other situations where individuals alter their environment.

Additionally, the late-life decline in resource stores has, unlike fat stores, consequences for generally unrelated individuals that did not physically interact with the focal individual: subsequent owners. The size of the old cone hoard of the current owner tended to be lower if they inherited a territory from a long-lived squirrel. This demonstrates it would be best to inherit a territory from an individual that died in the prime of their life, but this is also the age when red squirrels are least likely to die (Descamps *et al.* 2007). For the same reasons as described above for sex, we think red squirrel juveniles have limited ability to choose the territory they first settle at, and so would not be able to seek-out middens of prime-aged squirrels. Still, any factor that shortens individuals’ lives, and so causes individuals to die nearer the age of peak hoard size, should increase the number of cones passed onto to subsequent owners, the implications of which remain to be explored.

### Limited genetic effects on cone hoards, or indirect genetic effects of previous owner

Hoard sizes have been linked to survival and fitness (Larivée *et al.* 2010; LaMontagne *et al.* 2013; Haines 2017), yet we estimated very low direct additive genetic variance in this trait, and so it should not be able to respond to selection (indeed we see no trend across years). While genetic variation in caching behaviour must have originally existed to allow its evolution, that has likely now been eroded by consistent directional selection. Furthermore, given individuals do not possess genetic variance for the number of cones they place in the ground, it is not surprising we did not find any genetic variance in how they influence the parturition date or LRS of the next owner of the territory. Therefore, we do not need to consider IGEs from the previous owner when predicting how these traits might evolve. There was however a modest amount of variance for the IGE on old cone hoard size, although the credible intervals were very close to zero. Given the limited amount of direct genetic variance, the positive (but overlapping zero) DGE-IGE covariance is hard to interpret, but suggests that larger hoards could still evolve, if those with large hoards and high fitness leave the most cones for the next occupant. This is true for any trait which lacks direct genetic variance but is influence by IGEs such as maternal genetic effects (McFarlane *et al.* 2015), although indirect effects through extended phenotypes are generally not considered. Indirect effects through extended phenotypes may therefore provide a general route through which traits lacking direct genetic variance could evolve.

### Whose phenotype is it anyway?

The indirect effects from previous owners that we have identified have the potential to influence evolutionary processes. While we estimated no IGEs (nor any non-zero DGE-IGE covariances) for resource dependent traits and fitness, we still estimated phenotypic effects of previous owners on these traits. This means that evolutionary change will conform less well to models of evolution that only consider contemporary effects, as past individuals can influence current traits and fitness. Since stored resources in this system depend on the masting of spruce trees, past environments can influence contemporary resource dependent traits, requiring us to possibly model these ecological “memory” effects to understand trait distributions (Filotas *et al.* 2014). Maternal genetic effect models of evolution might be useful here, as although we have identified no IGEs (a maternal genetic effect is a type of IGE), such models do incorporate a lag in evolutionary change (Kirkpatrick & Lande 1989), something which may be occurring here. Quantifying how past environments, past individuals and historical selection can influence contemporary traits, and the consequences of this for ecological and evolutionary processes is an exciting next step for this line of research to take. In general, a given individual’s phenotype, and even its fitness, might only partly be under direct control, and some aspects of its phenotype may be influenced by individuals with whom it has not directly interacted through extended phenotypes (Dawkins 1982).

### Conclusions

Due to an extended phenotype, the food hoard size, the date of spring breeding, and the lifetime reproductive success of a red squirrel currently occupying a territory are influenced by the previous owner of the territory. This means the key traits, and even the fitness of an individual, are not under its direct control but are also influenced by previous individuals, and possibly previous environments. We did not find that cone hoard sizes were heritable, so in this case an extended phenotype did not give IGEs between individuals that did not meet, but this should be examined in other systems. A greater appreciation of what can be considered extended phenotypes, including the phenotypes of individuals when they interact with others, and their quantitative genetic underpinnings, will give us a greater appreciation of how these traits influence the ecology and evolution of populations.

## Acknowledgements

We thank Agnes MacDonald for long-term access to her trapline, and to the Champagne and Aishihik First Nations for allowing us to conduct work on their traditional territory. We thank all the volunteers, field assistants and graduate students whose tireless work makes the KRSP possible. We thank Andrea E. Wishart for useful comments on a previous draft. We have no conflicts of interest.

## Author contributions

DNF and AGM conceived of the research question. SB initiated the long-term study and all authors contributed to field logistics, data collection and the writing of the manuscript. DNF drafted the manuscript and conducted the data analysis, with guidance from AGM. All authors approved of the final manuscript for submission.

## Funding statement

Funding for this study was provided by the Natural Sciences and Engineering Research Council, the Northern Scientific Training Program, the National Science Foundation, and the Ontario Ministry of Research and Innovation

